# Selective Targeting of a Defined Subpopulation of Corticospinal Neurons using a Novel Klhl14-Cre Mouse Line Enables Molecular and Anatomical Investigations through Development into Maturity

**DOI:** 10.1101/2024.12.10.627648

**Authors:** Jake Lustig, Alexander Lammers, Julia Kaiser, Payal Patel, Aidan Raghu, James M. Conner, Phong Nguyen, Eiman Azim, Vibhu Sahni

**Affiliations:** Burke Neurological Institute, White Plains, New York 10605; Molecular Neurobiology Laboratory, Salk Institute for Biological Studies, La Jolla, CA, 92037; Feil Family Brain and Mind Research Institute, Weill Cornell Medicine, New York, New York 10065; Weill Cornell Graduate School of Medical Sciences, New York, NY, 10065

**Keywords:** Corticospinal, genomic Cre reporters, conditional AAV labeling, Crim1, Cerebellin, segmental axon projection, Brainstem innervation, Easi-CRISPR

## Abstract

The corticospinal tract (CST) facilitates skilled, precise movements, which necessitates that subcerebral projection neurons (SCPN) establish segmentally specific connectivity with brainstem and spinal circuits. Developmental molecular delineation enables prospective identification of corticospinal neurons (CSN) projecting to thoraco-lumbar spinal segments; however, it remains unclear whether other SCPN subpopulations in developing sensorimotor cortex can be prospectively identified in this manner. Such molecular tools could enable investigations of SCPN circuitry with precision and specificity. During development, Kelch-like 14 (*Klhl14*) is specifically expressed by a specific SCPN subpopulation, CSN_BC-lat_, that reside in lateral sensorimotor cortex with axonal projections exclusively to bulbar-cervical targets. In this study, we generated Klhl14-T2A-Cre knock-in mice to investigate SCPN that are *Klhl14+* during development into maturity. Using conditional anterograde and retrograde labeling, we find that Klhl14-Cre is specifically expressed by CSN_BC-lat_ only at specific developmental time points. We establish conditional viral labeling in Klhl14-T2A-Cre mice as a new approach to reliably investigate CSN_BC-lat_ axon targeting and confirm that this identifies known molecular regulators of CSN axon targeting. Therefore, Klhl14-T2A-Cre mice can be used as a novel tool for identifying molecular regulators of CST axon guidance in a relatively high-throughput manner *in vivo*. Finally, we demonstrate that intersectional viral labeling enables precise targeting of only Klhl14-Cre+ CSN_BC-lat_ in the adult central nervous system. Together, our results establish that developmental molecular delineation of SCPN subpopulations can be used to selectively and specifically investigate their development, as well as anatomical and functional organization into maturity.

**Significance Statement:** The cortex connects to brainstem and spinal targets through subcerebral projection neurons (SCPN), which exhibit molecular diversity during development based on their neocortical location and axonal targets. We generated a novel Klhl14-Cre mouse line to utilize this developmental delineation and drive Cre expression in a specific SCPN subpopulation. This developmental specificity enabled investigation of 1) areal locations of Klhl14+ SCPN in mature cortex, 2) their axonal collateralization at maturity, and 3) which genes can control their axon targeting. Using intersectional tools, we can also selectively label these neurons in the adult CNS. Therefore, developmental molecular delineation of SCPN not only provides prospective identification but also enables molecular analysis during development, as well as anatomical and functional investigations in adulthood.

## Introduction

The corticospinal tract (CST) is a principal circuit responsible for skilled voluntary movements (Martin, 2005; Sahni et al., 2020; Welniarz et al., 2016). For such skilled motor control, it is critical that corticospinal neurons (CSN), which form a subset of all subcerebral projection neurons (SCPN), establish appropriate connectivity with their segmentally distinct targets. This necessitates that cortico-brainstem neurons project exclusively to the brainstem, while corticospinal projections extend to segmentally appropriate spinal targets in the cervical, thoracic, and lumbar cord. This segmental targeting specificity arises, in part, via axon extension specificity during development that is maintained into maturity (Sahni et al., 2021b). CSN subpopulations residing in distinct neocortical locations project to segmentally distinct targets (Sahni *et al*., 2021b). CSN in lateral cortex project axons exclusively to targets in the brainstem and cervical cord (CSN_BC-lat_). CSN in medial cortex (CSN_med_) are relatively more heterogeneous– a subpopulation extends axons exclusively to the brainstem and cervical cord (CSN_BC-med_) and resides interdigitated with another subpopulation that extends axons past these proximal targets to thoraco-lumbar segments (CSN_TL_).

These anatomically distinct subpopulations are also molecularly distinct during development (Sahni *et al*., 2021b). *Kelchlike-14* (*Klhl14*) is specifically expressed by CSN_BC-lat_, while *Cysteine rich transmembrane BMP regulator 1* (*Crim1*) is specifically expressed by CSN_TL_. We previously tested the hypothesis that developmental molecular delineation could prospectively identify these subpopulations into maturity. Using an inducible Crim1CreERT2 mouse line, we established that *Crim1* expression during development prospectively identified CSN_TL_ even before their differential axon targeting from other CSN subpopulations was evident. However, whether this developmental molecular delineation could enable identification and manipulation of other CSN subpopulations at maturity remained unknown. We therefore investigated whether CSN_BC-lat_ could similarly be identified during development via the expression of *Klhl14*. We generated Klhl14-T2A-Cre knock-in mice to label *Klhl14*+ SCPN during development and investigate their axonal projections at maturity. Our hypothesis was that SCPN, labeled via *Klhl14* expression during development, would only extend axons to bulbar-cervical segments at maturity. Surprisingly, however, we find that when we breed Klhl14-T2A-Cre mice with genomic Cre reporter mice, labeled SCPN axons extend to all levels of the neuraxis. Since this strategy labels all neurons that express *Klhl14* at any time during development, we next used a different approach to specifically label *Klhl14*+ SCPN at the developmental times when we had previously confirmed *Klhl14* expression specificity. Using conditional viral labeling, we find that Cre expression in Klhl14-T2A-Cre mice faithfully recapitulates this previously established specificity of *Klhl14* expression. Klhl14-Cre+ SCPN labeled at P0 reside in lateral sensorimotor cortex and extend axons exclusively to brainstem and cervical segments i.e., *Klhl14* expression prospectively delineates CSN_BC-lat_. By comparing the axonal projections of *Klhl14*+ SCPN with all SCPN in lateral sensorimotor cortex, we establish that Cre expression in Klhl14-Cre mice labels nearly all CSN_BC-lat_. Our results highlight a limitation of using genomic Cre reporter mice when taking advantage of developmental molecular delineation to investigate CSN subpopulations into maturity. Given the highly dynamic nature of gene expression during development, this limitation is likely applicable to other neuronal populations beyond CSN, indicating that genomic Cre reporters alone may be insufficient for investigating such parcellation.

We also establish the use of Klhl14-T2A-Cre mice as a tool to identify genes regulating CSN axon extension specificity during development. Our previous work had identified that misexpression of Crim1 (Sahni et al., 2021a) and Cbln1 (Song et al., 2023) in lateral sensorimotor cortex is sufficient to redirect CSN_BC-lat_ axons past their targets in the brainstem and cervical cord toward thoraco-lumbar spinal segments. In these investigations, we solely relied on the anatomical separation of CSN_BC-lat_ from CSN_med_. Klhl14-T2A-Cre mice enable selective labeling of CSN_BC-lat_ even in instances where AAV injections spread into medial cortex. Molecular delineation therefore provides a significant advance over using anatomical separation alone to target and investigate distinct CSN subpopulations. Finally, we describe the use of an intersectional viral labeling approach combining both Cre and Flp-dependent recombination to specifically target *Klhl14*+ CSN_BC-lat_ into maturity. Together, these results show that Cre expression in Klhl14-T2A-Cre at P0 labels all CSN_BC-lat_. More broadly, our results indicate that early developmental molecular delineation of distinct CSN subpopulations can be used to specifically label them prospectively with reliability and precision. These approaches can in turn enable investigating their anatomy and circuit-level function into maturity.

## Materials and Methods

### Generation of Klhl14-Cre Mice

Klhl14-3xHA-T2A-iCre mice were generated to carry three HA-tags inserted at the C-terminus of Klhl14 protein followed by a T2A-iCre sequence. These mice were generated in the Transgenic Core Facility at the Salk Institute for Biological Studies, La Jolla, CA using the Easi-CRISPR strategy (Quadros et al., 2017). For this, long single-stranded DNA donors (ssDNAs, HDR template containing 3x HA tags-T2A-iCre flanked by 2 homology arms, 1634 bp) were ordered from IDT (Megamer™). These were injected into one-cell staged mouse zygotes along with two sgRNAs (made by Synthego) and Cas9 ribonucleoprotein (obtained from IDT) that together formed the CRISPR RNPs complex. The following two sgRNAs were used 1) Klhl14-sgRNA1: TGGATGGTGGCACTATGCCG; and 2) Klhl14-sgRNA2: ACCCTACAACAAATGACAGC.

For the microinjections, 3-4 weeks old C57BL/6J donor female mice (Jackson Lab) were superovulated by 5 IU of PMSG and 5 IU of HCG injections and mated 1:1 with males. The next morning, one-cell stage embryos were collected and cultured in microdrops of KSOM+AA media (Millipore) covered with embryo-tested mineral oil at 37C in 5% CO2. CRISPR RNP mixture was freshly prepared before microinjection by resuspending Cas9 protein (IDT), two sgRNAs (Synthego) and ssDNA (IDT) in microinjection buffer (10 mM Tris, pH 7.4–7.5, 0.15 mM EDTA) at a concentration of 20:10:10:5 ng/μl. CRISPR mix was injected into the male pronuclei using a Nikon Eclipse microscope and Narishige micromanipulators. Injected embryos were surgically transferred into the oviducts of 0.5 days post coitus (dpc) pseudopregnant CD1 females (Charles River).

To confirm the correct insertion of the 3xHA-T2A-iCre into the Klhl14 locus, genomic DNA from the putative targeted pups were used for genotyping. The following primers were used to amplify the targeted insertion, which were located outside the ssDNAs (HDR template) in the Klhl14 locus: 1) Klhl14-gt-5a: GCAGAGGTCTTCCCAGTGTAGC; and 2) Klhl14-gt-3a: AGTTTGTGTAAAGGCGGAACAAAGC. The resulting 1703bp PCR product was cloned and sequenced to confirm that it was correctly targeted into the Klhl14 locus. Targeted pups were bred with the wild-type mice to generate the first generation (F1) of the heterozygous mice. Once the Klhl14-3xHA-T2A-iCre mouse line was established, the following primers were used in a multiplex reaction for genotyping and to distinguish between Klhl14-Cre WT, Heterozygous and homozygous mice:

<u>Klhl14-Cre</u> (common forward):

GAACCTGGACAGAACTCGA

<u>Klhl14-WT Reverse</u>:

AGCCTGATATGGAAGAGTCTG

<u>Klhl14-Cre Reverse</u>:

ACACAGACAGGAGCATCTTC

This reaction results in bands of the following sizes: WT allele-334 bp; Klhl14-Cre allele-467 bp.

### Mice

To generate mice for experimental analyses, we only used heterozygous Klhl14-Cre mice. For this, male Klhl4-Cre homozygous mice were mated with female CD-1 WT mice (Charles River Laboratories, Wilmington, MA) and the resulting pups were used for both anterograde injections and retrograde injections. P0 was set as the day of birth. Both male and female mice were used in all analyses. The transgenic mouse used for Figure 1E resulted from mating a male Klhl14-Cre heterozygous mouse with female Emx1FlpO;Ai65 heterozygous mice. Emx1FlpO and Ai65 mice were maintained and genotyped as previously described (Madisen et al., 2015; Sahni *et al*., 2021b). Mice received food and water ad libitum and were housed on a 12-hour on/off light cycle. All mouse studies were approved by the IACUC at Weill Cornell Medicine and the Salk Institute for Biological Studies, and all studies were performed in accordance with institutional and federal guidelines.

**Figure 1.**
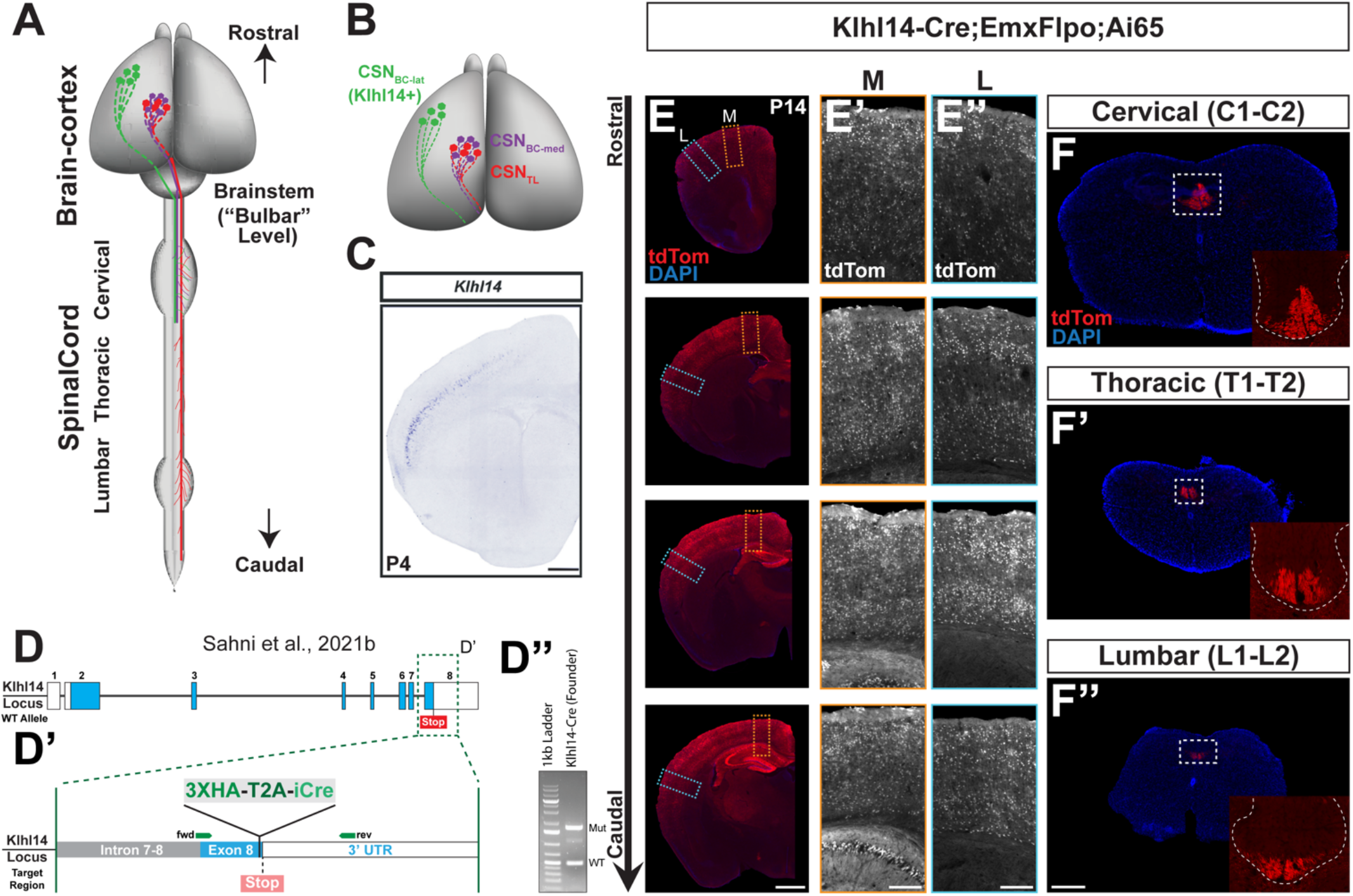
Genomic reporters of Cre-dependent recombination do not identify SCPN subpopulation specificity of Cre expression in Klhl14-Cre mice. ***A-B***, Schematic (adapted from Sahni et al., 2021b) showing the locations of distinct CSN subpopulations and their corresponding axonal projections in mouse CNS. CSN_BC-lat_ (green) reside in rostrolateral cortex and extend axons only to bulbar-cervical targets. CSN_BC-med_ (purple) reside in medial cortex and extend axons only to bulbar-cervical targets. CSN_TL_ (red) reside in medial cortex and extend to thoraco-lumbar levels, but extend collaterals across all spinal levels. *Klhl14* is only expressed by CSN_BC-lat_. ***C***, In situ hybridization image on a coronal section of a P4 mouse brain showing *Klhl14* expression by CSN_BC-lat_ (from Sahni et. al., 2021b). ***D***, Schematic showing the genomic organization of the WT Klhl14 locus. Klhl14 is encoded by 8 exons. (D’) Magnified view of region boxed in D showing the point of insertion of Cre in Klhl14-Cre mice. The T2A-Cre sequence was targeted to exon 8 immediately 3’ to the STOP codon. The remaining part of the 3’ UTR in exon 8 was used as a homology arm. (D”) Genomic PCR from F2 mice confirmed proper insertion of Cre, with a resultant product of 1703 bp. ***E***, Coronal brain sections collected from a Klhl14-Cre;Emxflpo;Ai65 transgenic mouse. Klhl14-Cre positive cell bodies are labeled via tdTomato reporter expression (in red and in monochrome). TdTomato+ neurons span all layers across both medial (E’) and lateral (E”) cortex. **F**, Axial spinal sections from the same Klhl14-Cre;Emxflpo;Ai65 transgenic mouse taken at cervical, thoracic, and lumbar levels. TdTomato+ axons are present in the dorsal funiculus at all spinal levels consistent with the widespread recombination in cortex. Scale bars 500 µm for **C**, 1 mm for **E**, and 250 µm for **E’-E”**, and **F**.

### Generation of AAV Particles

The source, titers, and additional details for all AAVs used in this study are listed in the table below.

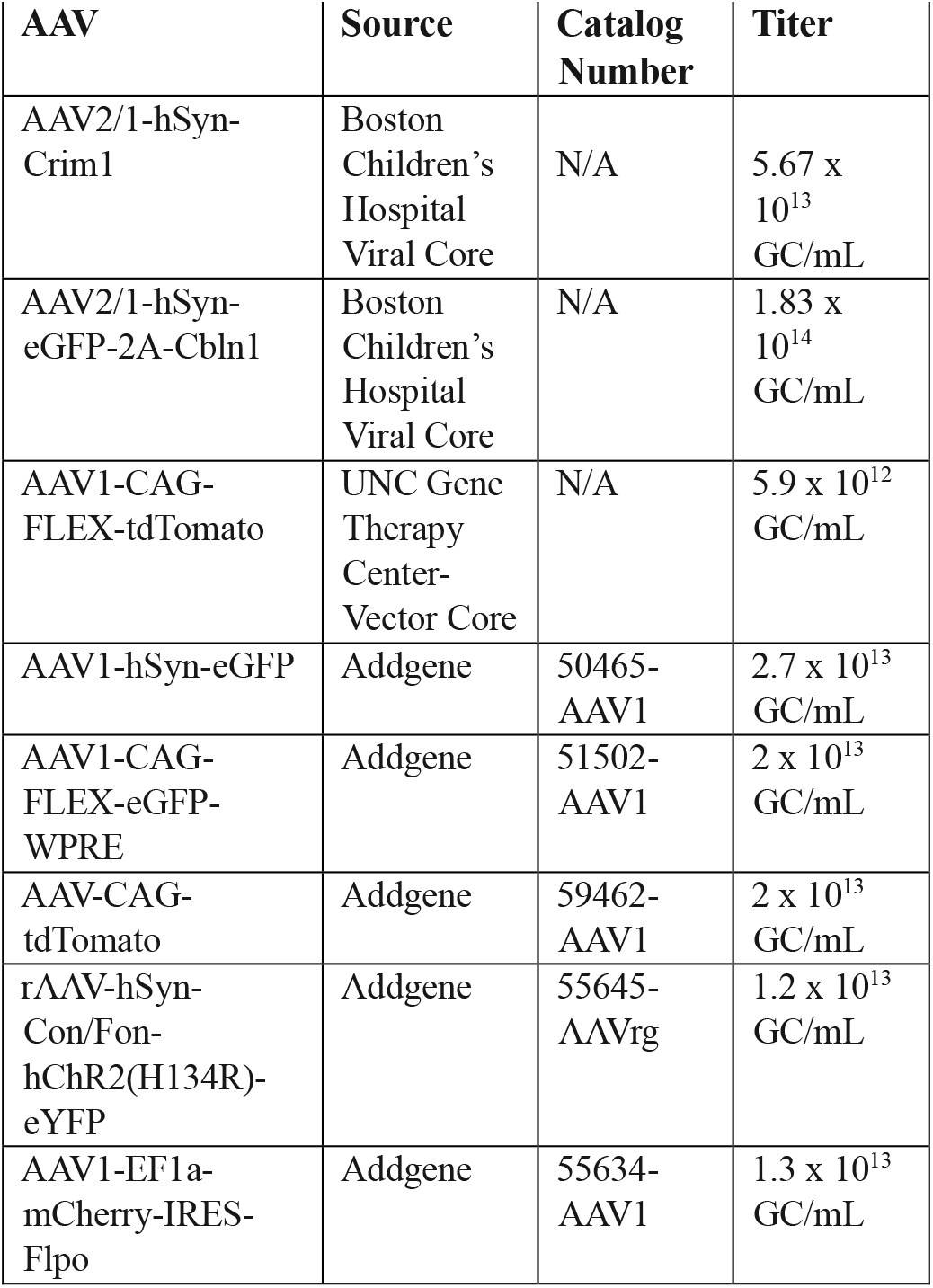

Constitutive and Cre-dependent AAV reporters were obtained either from Addgene (Watertown, MA) or from the University of North Carolina Gene Therapy Center Vector Core.

For generating AAVs to overexpress Crim1 and Cbln1, both genes were cloned into shuttle plasmids, where gene overexpression is driven by the human synapsin promoter (hSyn). For Cbln1, an eGFP-T2A coding sequence was placed in frame 3’ to the Cbln1 ORF. For AAV-Crim1, a GFP reporter could not be introduced since this exceeds the size limitations for AAV packaging. AAV particles were packaged by the Boston Children’s Hospital Viral Vector Core.

### Ultrasound guided intracranial and intraspinal AAV Injections

All AAV injections were performed in postnatal mice under visual guidance provided by ultrasound-mediated backscatter microscopy (Vevo 2100; VisualSonics, Toronto, ON, Canada) using previously described protocols (Sahni *et al*., 2021a; Sahni *et al*., 2021b; Song *et al*., 2023). For all injections, mouse pups were anesthetized using hypothermia and injections were performed using a beveled glass micropipette attached to a nanojector (Nanoject III; Drummond Scientific, Broomall, PA). Following injections, pups were then placed on a heating pad and returned to their dams in home cages.

For intracortical injections for anterograde labeling, the micropipette was inserted unilaterally into rostrolateral cortex using defined landmarks. Mice received 2 sets of 15 injections of 23nL each: one each at 2 different rostrocaudal levels. Injections were performed at 1-second intervals, with a 10-second waiting period after the 15^th^ injection before the micropipette was removed. Retrograde labeling from the cerebral peduncles was performed at P0, while labeling from the cervical cord was performed at P2 using established protocols (Sahni *et al*., 2021a; Sahni *et al*., 2021b; Song *et al*., 2023). For labeling from the cerebral peduncle, AAV injections were delivered in the rostral pons as a set of 7 injections of 23nL each, at 3 different injection sites per hemisphere. For labeling from the cervical cord, we injected 10 injections of 23nL each on either side of the midline.

For intersectional viral labeling in Figure 6, mice were either injected at P3 with rAAV-FLEX-eGFP or at P2 with rAAV-Cre-On-Flp-On-eYFP bilaterally into the cerebral peduncle. On the same day, mice which received the Cre-On-Flp-On-eYFP also received intracortical injections with AAV-mCherry-Flpo into rostrolateral cortex. For these injections, mice received 1 set of 15 injections (per cerebral hemisphere) of AAV-mCherry-Flpo at 23nL per injection.

### Tissue collection and preparation

P14 or P15 mice were euthanized using an intraperitoneal injection of 150 mg/kg ketamine along with 15 mg/kg xylazine. Mice were transcardially perfused, first with 1x PBS, followed by 4% paraformaldehyde (PFA). The brain and spinal cord were dissected and postfixed in 4% PFA overnight at 4ºC. Samples were rinsed with 1x PBS and stored in PBS-azide (0.025% sodium azide in 1x PBS) at 4ºC for long-term storage. For downstream processing, samples were first cryoprotected in sucrose PBS (30% sucrose in 1x PBS), frozen in Tissue-Tek OCT compound (Sakura Finetek, Torrance, CA), and cryosectioned on a Leica CM3050 S cryostat (Leica microsystems, Germany). For analyzing the cortex, prior to freezing, the cerebellum, pons, and medulla were removed as one tissue block from the forebrain using a razor blade. 50 μm coronal brain sections were collected in PBS-azide. The brainstem samples were then frozen separately and serial sections collected in PBS-azide. For axon quantification analysis on axial spinal sections, 50 µm axial sections from C1-C2 and T1-T2 spinal cord were collected in PBS-azide. For horizontal sections shown in Figure 5, the thoracic cord was separated from the lumbar cord at T13/L1 using a razor blade. An orientation cut was made at the caudal end prior to freezing to identify the rostral and caudal ends of the cord in horizontal sections. 70 µm thick horizontal sections were collected serially in 1X PBS.

### Immunohistochemistry

Immunohistochemistry was performed as previously described (Sahni *et al*., 2021a; Sahni *et al*., 2021b; Song *et al*., 2023). Sections were blocked for 30 minutes at room temperature in 1x PBS containing 0.3%BSA and 0.3% Triton X 100, followed by overnight incubation at 4ºC with the primary antibody, which was diluted in the same blocking solution. Following 3 rinses in 1x PBS, sections were incubated for 3 hours at room temperature in secondary antibodies (1:750) in the same blocking solution. After 3 rinses in 1xPBS, sections were mounted onto glass slides. Cover slips were affixed using DAPI Fluoromount-G (SouthernBiotech, 0100-20, Birmingham, AL) and sealed using clear nail polish. Slides were then allowed to air-dry overnight before being imaged. The following primary antibodies were used: Rabbit-anti-RFP (1:750; Rockland Immunochemicals, Pottstown, PA) for analyzing tdTomato+ axons in the brainstem and spinal cord, Rabbit anti-GFP (1:500; ThermoFisher Scientific, A-11122, Waltham, MA) for analyzing both eGFP and YFP fluorescence in the brainstem in Figure 6. The following secondary antibodies were used: AlexaFluor 546 goat-anti-rabbit (ThermoFisher Scientific, A-11035), AlexaFluor 488 goat-anti-rabbit (Thermofisher Scientific, A-11008). Coronal forebrain (cortical) sections did not undergo immunohistochemical processing. Free-floating sections were mounted on glass slides and cover slipped as above.

### Microscopy and Imaging

Prior to tissue sectioning, all brains were first whole mount imaged (dorsal view) on a Nikon SMZ18 stereomicroscope (Nikon Japan). Epifluorescence images were acquired on a Zeiss Axio Imager M2 microscope (Zeiss, Oberkochen, Germany) using Stereo Investigator Software (MBF Biosciences, Williston, VT). For axon extension quantification on axial spinal sections, labeled axons in the dorsal funiculus were imaged on a Leica SP8 confocal microscope (Leica Microsystems, Wetzlar, Germany) with 63x oil immersion lens using LASX software (Leica Microsystems). Confocal images of horizontal sections of the thoracic cord were acquired at 20x and maximum intensity projections were produced using Fiji ImageJ (National Institutes of Health, Bethesda, MD).

### 3-Dimensional Injection Volume Reconstruction

To create 3-dimensional (3D) model representations of the injected cortical volumes of tdTomato+ and eGFP+ neurons, we utilized “VOL3D” (designed by Dr. Julia Kaiser; available on GitHub: https://github.com/jkaiser87/VOL3D, 2024) to transfer coordinates from 2D brain slices into the Common Coordinate Framework version 3 brain (CCFv3, 10 um atlas) (Wang et al., 2020). Briefly, the injection volumes of eGFP+ and tdTomato+ neurons in coronal brain slices were outlined in FIJI/ImageJ (version 1.54f, (Schindelin et al., 2012)). Coordinates of the outlines were transferred into CCFv3 space through affine transformations using our custom-made MATLAB (MathWorks, Inc., Natick, MA) script, largely based on “AP_histology” (https://github.com/petersaj/AP_histology, 2019/2024) [Source Code](Peters, 2024). CCFv3-adapted coordinates were used to compute surface boundaries via Delaunay triangulation, followed by Laplacian smoothing to reconstruct and display the 3D injection volumes within the CCFv3 framework.

### Quantification and spatial representation of retrogradely labeled cells

Retrogradely labeled Klhl14-Cre+ SCPN/CSN were detected using AMaSiNe (Song et al., 2020) following the recommended protocol (https://github.com/vsnnlab/AMaSiNe, 2020/2021 (Song et al., 2020)) with minor manual edits to obtain a 3D representation of the location of traced neurons. Quantification of numbers of retrogradely labeled neurons was generated using NeuroInfo (MBF BioSciences, Williston, VT).

### Axon extension Quantification

For axon extension analysis in the spinal cord, we only included mice in which there was detectable tdTomato fluorescence in the whole mount images. For quantification, confocal images were analyzed using a custom-made macro in Fiji. Briefly, all confocal Z-stacks were merged to create a maximum projection image. The dorsal funiculus was manually annotated as a region of interest (ROI) in this image. The ROI was thresholded in the 3 brightest Z-planes to create a binary image, and the area of thresholded pixels was summed. For each animal, 3 axial sections were analyzed at each spinal level (cervical and thoracic) and the area averaged. For each animal, the average area at thoracic T1-T2 was divided by the average area at cervical C1-C2 to obtain the T:C ratio, which was expressed as a percentage. Data are presented as mean ± SEM (standard error of mean), with *n* indicating the number of mice used in each group for comparison. Male and female mice were used without distinction in experiments.

## Results

### Genomic Cre-reporter mice show that Klhl14-T2A-Cre drives widespread recombination across the neocortex

Our previous work has established that CSN in lateral cortex, i.e. CSN_BC-lat_ are relatively more homogeneous than CSN in medial cortex, whereby CSN_BC-lat_ extend axons exclusively to bulbar-cervical segments (schematized in Figures 1A, B) (Sahni et al., 2021b). We had established *Klhl14* expression using *in situ* hybridization. In the developing neocortex, from E18.5 to P7, *Klhl14* is expressed by SCPN in lateral, but not medial, layer V, i.e., *Klhl14* is specifically expressed by CSN_BC-lat_ (Figure 1C). However, these analyses could not investigate the axonal trajectory of *Klhl14*-expressing neurons during development at maturity. Therefore, to establish this, we generated Klhl14-T2A-Cre knock in reporter mice (hereby referred to as “Klhl14-Cre”), in which we introduced a 3xHA-T2A-iCre cassette in frame with the Klhl14 coding sequence immediately 5’ to the STOP codon (Figure 1D, D’). Proper insertion of the Cre coding sequence within the Klhl14 locus was confirmed by PCR (Figure 1D”) followed by sequencing.

We next used these mice to investigate axonal projections of *Klhl14*+ SCPN. We crossed Klhl14-Cre mice with Emx-IRES-FlpO (Sahni *et al*., 2021b) and Ai65(Madisen *et al*., 2015) reporter mice, to generate Klhl14T2A-Cre; Emx1-IRES-FlpO; Ai65 triple transgenic mice. Since *Klhl14* is expressed by a subpopulation of spinal interneurons, we used Emx1-IRES-FlpO, which drives FlpO recombinase expression exclusively in neocortical projection neurons, with no spinal expression. In Ai65 reporter mice, tdTomato expression only occurs upon both Cre- and FlpO-mediated recombination. This breeding strategy therefore enabled us to avoid labeling any spinal *Klhl14*-expressing cells and specifically investigate tdTomato+ axonal projections of *Klhl14*+ SCPN. Our prediction was that only CSN_BC-lat_ would be tdTomato+ in these mice. However, we find that tdTomato+ neurons span all cortical layers and not just layer V, and also reside across the entire medio-lateral axis (Figure 1E-E”). Furthermore, and in line with the distribution of tdTomato+ neurons in cortex, we find that tdTomato+ SCPN axons extend beyond the cervical cord to thoracic and lumbar levels (Figure 1F-F”). These results would suggest that Cre expression in Klhl14-Cre mice does not recapitulate the known expression of *Klhl14*. However, because this breeding strategy with genomic Cre-reporter mice does not allow for temporal control over the timing of recombination, it remained possible that Cre expression in Khl14-Cre mice occurred correctly at the appropriate developmental stages. We therefore tested this possibility via conditional anterograde and retrograde labeling at these developmental stages using AAV reporters.

### Conditional retrograde labeling identifies specificity of Cre expression by CSN_BC-lat_ and cortical location of CSN_BC-lat_ at maturity

One possible explanation for the broad Cre-reporter expression when using genomic Cre reporter mice is that *Klhl14* is expressed more broadly, prior to P0, in the embryonic neocortex. This would in turn lead to more widespread Cre-mediated recombination across sensorimotor cortex. Another possibility is that Cre expression in Klhl14-Cre mice is being driven by other genomic regulatory elements and therefore, as a result, Cre expression does not reflect *Klhl14* expression. This latter possibility would indicate Klhl14-Cre mice cannot be used to investigate any *Klhl14*+ neuronal populations. To distinguish between these two possibilities, we used retrograde AAVs (rAAV) (Tervo et al., 2016) to conditionally label all Cre+ SCPN in Klhl14-Cre mice at P0, i.e., past embryonic developmental stages and at a time when we had previously identified Klhl14 expression specificity (Sahni *et al*., 2021a; Sahni *et al*., 2021b).

For this, we co-injected two rAAVs into the cerebral peduncle of Klhl14-Cre mice at P3: one constitutively expressing tdTomato (rAAV-CAG-tdTomato) and one that conditionally expressed eGFP upon Cre expression (rAAV-FLEX-eGFP) (Figure 2A). If Cre expression in Klhl14-Cre mice faithfully reflected our previously established *Klhl14* expression (as shown in Figure 1C)(Sahni *et al*., 2021a; Sahni *et al*., 2021b), we would predict that eGFP+ SCPN would be restricted to lateral but not medial cortex. However, if Cre expression does not reflect *Klhl14*, then eGFP+ SCPN would be present across both medial and lateral sensorimotor cortex, similar to our results obtained using genomic Cre reporter mice shown in Figure 1E. We analyzed injected mice at P14. We find that while SCPN across both medial and lateral cortex are tdTomato+, eGFP+ SCPN are restricted to lateral sensorimotor cortex (Figure 2A’, A”). We also performed 3D AMaSiNe reconstructions of eGFP+ SCPN across 3 different Klhl14-Cre mice and find that all *Klhl14*+ SCPN reside in lateral sensorimotor cortex (pink in Figure 2C). Together, these results confirm that eGFP+ SCPN are CSN_BC-lat_. This further indicates that Cre expression in Klhl14-Cre mice does reflect *Klhl14* expression and that the widespread recombination seen when using genomic Cre reporter mice likely reflects broader *Khl14* expression in embryonic neocortex.

**Figure 2.**
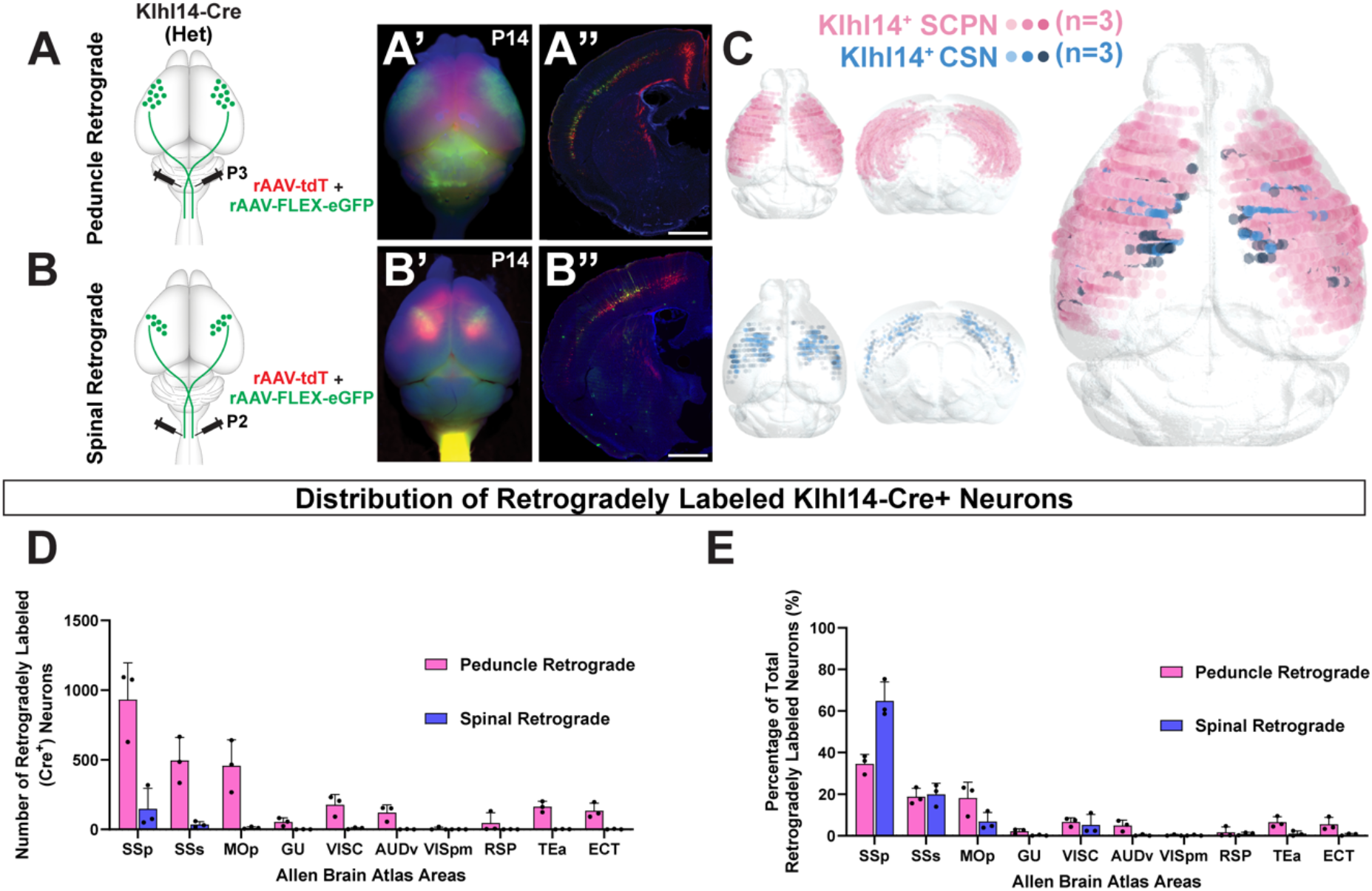
Conditional retrograde labeling identifies specificity of Klhl14-Cre expression by CSN_BC-lat_ enabling identification of their cortical location at maturity. ***A-A”***, Klhl14-Cre heterozygous mice were co-injected with rAAV-FLEX-eGFP and rAAV-tdTomato in the cerebral peduncle at P3 (schematized in A) and the brains collected at P14. ***A’***, Wholemount imaging of the injected brain finds retrogradely labeled tdTomato^+^ SCPN span the entire sensorimotor cortex while eGFP^+^ (Klhl14-Cre+) SCPN occupy a smaller area of cortex, specifically residing in lateral cortex. ***A’***, Coronal sections of the same brain reveal that eGFP^+^ SCPN reside laterally (where Klhl14^+^ SCPN are known to reside), while tdTomato^+^ SCPN reside in both medial and lateral cortex. ***B-B”***, Klhl14-Cre heterozygous mice were co-injected with rAAV-FLEX-eGFP and rAAV-tdTomato in cervical spinal cord at P2 (schematized in B) and brains collected at P14. ***B’***, Wholemount imaging finds fewer tdTomato+ CSN labeled than SCPN in A’. eGFP^+^ CSN occupy a smaller area of cortex than tdTomato^+^ CSN, indicating specificity of Cre expression. ***B”***, Coronal sections of the same brain find that while tdTomato^+^ CSN span both medial and lateral cortex, eGFP^+^ CSN reside laterally (where Klhl14^+^ SCPN are known to reside). Further, there are fewer eGFP^+^ CSN than GFP+ SCPN labeled in A”. ***C***, 3D AMaSiNe reconstructions of conditionally labeled neurons from 3 Klhl14-Cre mice each that received retrograde labeling from either the cerebral peduncle (Klhl14-Cre+ SCPN, pink) or cervical spinal cord (Klhl14-Cre+ CSN, blue). SCPN occupy a much larger area of cortex compared to CSN, but both primarily reside in lateral cortex. ***D***, Distribution of Klhl14^+^ SCPN across different cortical areas (annotations as assigned by the Allen Brain Atlas). Most Klhl14-Cre+ neurons reside in somatosensory cortices. ***E***, Percentage counts of labeled Klhl14-Cre+ neurons in each area, again demonstrating that the majority of Klhl14+ SCPN and CSN reside in sensory cortices. Scale bars are 1mm.

Our recent results have found that there is additional diversity within CSN_BC-lat_ and that they comprise at least two additional subpopulations: 1) a subpopulation that only extends axons to the brainstem (CBN); and 2) a subpopulation of cervical-projecting CSN (CSNc), which reside interdigitated with CBN (Kaiser et al., 2022). We therefore investigated whether Klhl14-Cre is expressed by one or both these CSN_BC-lat_, i.e., whether Klhl14-Cre only labels a subset of CSN_BC-lat_. To test this, we first asked whether Klhl14-Cre+ SCPN extend axons to the cervical spinal cord. If Klhl14-Cre is only expressed by CBN, conditional retrograde labeling from the cervical cord would not label any Klhl14-Cre SCPN. We therefore co-injected the identical combination of rAAVs as in Figure 2A from the cervical spinal cord at P2 (schematized in Figure 2B). Since CSN form a subset of SCPN, we find that overall, rAAV-CAG-tdTomato injection from the cervical cord labels fewer neurons than injections from the cerebral peduncle (compare Figure 2A’ and B’). Importantly, however, we do find eGFP+ CSN labeled in Klhl14-Cre+ mice indicating that spinal-projecting neurons express Klhl14-Cre (Figure 2B’, B”). 3D AMaSiNe reconstructions of eGFP-labeled CSN across 3 different Klhl14-Cre mice show Klhl14-Cre CSN reside in lateral sensorimotor cortex (blue neurons in Figure 2C). Interestingly, these reconstructions show that Klhl14-Cre+ CSN occupy a smaller area in lateral cortex as compared to Klhl14-Cre+ SCPN. This suggested that Klhl14-Cre is also expressed by a subpopulation of CSN_BC-lat_ that do not extend axons to the cervical spinal cord, i.e. CBN. To address this question, we quantified the number of retrogradely labeled Klhl14-Cre+ SCPN vs. Klhl14-Cre+ CSN (Figure 2D, E). We find that there are significantly higher numbers of neurons retrogradely labeled from the cerebral peduncle than the cervical spinal cord (Figure 2D, E). This indicates that Klhl14-Cre is also expressed by neurons that extend axons to the cerebral peduncle, but do not extend axons to the cervical spinal cord, i.e. CBN. Collectively, these quantitative results show that within the broad population of CSN_BC-lat_, Klhl14-Cre is expressed by both CBN and cervical-projecting CSN in lateral sensorimotor cortex.

Since our previous developmental *Klhl14* expression analyses were performed in the postnatal cortex (Sahni et al., 2021a and 2021b), we could not accurately map areal locations of *Klhl14*+ SCPN within the cortex, since distinct cortical areas have not been fully defined at these early postnatal ages. Using injected Klhl14-Cre mice, we now had the ability to not only quantify the number of labeled neurons, but also their relative distribution across different cortical areas in the P14 mouse cortex. Consistent with our previous expression analyses that identified *Klhl14* is expressed in lateral sensorimotor cortex, we find that *Klhl14*+ SCPN largely reside in somatosensory cortical areas (Figure 2D, E). Together, these data reveal that Klhl14-Cre faithfully recapitulates our previously established *Klhl14* expression by CSN_BC-lat_ and that *Klh1l4*+ is expressed by both CBN and CSN in lateral sensorimotor cortex.

### Conditional anterograde labeling in Klhl14-T2A-Cre mice confirms that Klhl14+ CSN do not extend axons to the thoracic cord

Our prior work had confirmed that CSN_BC-lat_ do not extend axons beyond the cervical spinal cord to thoraco-lumbar levels (Sahni et al., 2021b). Given that *Klhl14* is expressed by CSN in lateral sensorimotor cortex, we next sought to investigate whether *Klhl14*+ CSN maintain the same axon targeting specificity in Klhl14-T2A-Cre mice. We therefore performed conditional anterograde labeling from lateral sensorimotor cortex. We co-injected at P0, the lateral cortex of Klhl14-Cre mice with two AAVs – 1) an AAV that constitutively expressed eGFP (AAV-hsyn-eGFP); and 2) an AAV that conditionally expressed tdTomato upon Cre-mediated recombination (AAV-FLEX-tdTomato). Brain and spinal cords from these injected mice were analyzed at P15 (strategy schematized in Figure 3A). We first investigated expression of both reporters in cortex. A series of coronal brain sections from one such injected mouse is shown in Figure 3B. We find that while the constitutive eGFP reporter expression spans much more broadly across lateral sensorimotor cortex, tdTomato expression remains confined to deeper layers. Even though the injections were performed in lateral cortex, there is notable spread of eGFP (non-Cre dependent reporter) expression into medial cortex (Figure 3D, D”). However, in striking contrast, tdTomato (Cre-dependent reporter) expression remains confined to lateral cortex (Figure 3D, D’) indicating specificity of Klhl14-Cre expression to CSN_BC-lat_. 3D reconstruction of the overall cortical territory occupied by labeled neurons from the same injected mouse show that while eGFP+ neurons extend into medial cortex, tdTomato+ (Cre-reporter) neurons remain confined to lateral cortex (Figure 3C).

**Figure 3.**
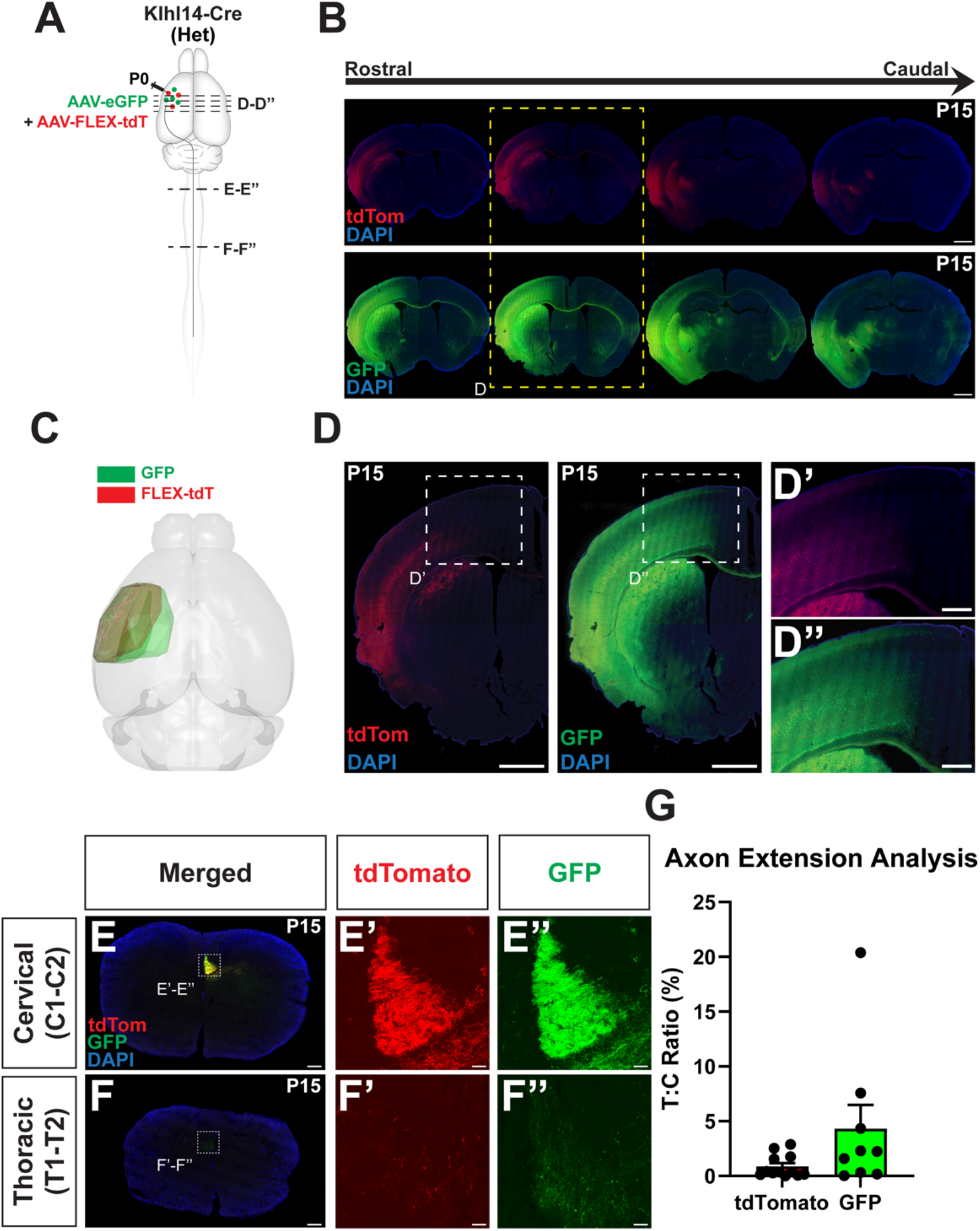
Anterograde Labeling in Klhl14-T2A-Cre Mice Confirms that Klhl14+ CSN do not extend axons to the Thoracic Cord. **A**, Schematic of experimental outline: Klhl14-Cre heterozygous mice were co-injected with AAV-eGFP and AAV-FLEX-tdTomato at P0 in rostrolateral cortex, where Klhl14-Cre+ SCPN reside. Samples were collected at P15. **B**, Series of coronal brain sections from one injected mouse spanning rostral (left) to caudal (right) showing localization and spread of both AAVs within cortex. **C**, Sample 3D reconstruction of an injected brain displaying the injection locations and volumes for both eGFP and FLEX-tdTomato. Note that the eGFP (non-Cre dependent) reporter expression spans more medially than tdTomato (Cre-dependent). **D**, Zoomed in images for the section marked by dotted yellow outline in (B). AAV-FLEX-tdTomato only infects cells in lateral cortex, where Klhl14-Cre SCPN reside (D’). Unlike AAV-FLEX-tdTomato, AAV-eGFP infects cells in both lateral and medial cortex (D”). **E-F”**, Spinal axial sections from an injected Klhl14-Cre mouse at cervical (E-E”) and thoracic (F-F”) levels. While both tdTomato+ (E’) and eGFP+ axons (E”) are seen in the cervical dorsal funiculus, tdTomato+ axons (F’) are not present in thoracic spinal cord. AAV-eGFP labeled axons (F”) are labeled in the thoracic spinal cord, likely due to nonspecific labeling of CSN_med_ populations. **G**, Quantification of thoracic to cervical ratio (T:C ratio) from injected Klhl14-Cre mice at P15. T:C ratio is the percentage of labeled axons in the cervical dorsal funiculus that extend to the thoracic cord. (n=12 for tdTomato+ axons; n=9 mice for GFP+axons). There is much more variability in the T:C ratio for eGFP+ axons than tdTomato+ axons. Scale bars are 1mm for **B, D**, 100 µm for **D’-D”**, 250 µm for **E, F** and 20 µm for insets in **E’-E”, F’-F”**.

We next analyzed axon extension in the spinal cord by comparing eGFP+ vs. tdTomato+ CSN axons in axial spinal sections from injected Klhl14-Cre mice. We find both eGFP+ and tdTomato+ axons are present in the cervical dorsal funiculus confirming that Klhl14-Cre is expressed by spinally-projecting neurons in lateral sensorimotor cortex (Figure 3E, E’, E”). While some eGFP+ axons can be seen in the thoracic dorsal funiculus, likely originating from the medial cortex, i.e., CSN_med_ (as shown in Figure 3D”), the majority of tdTomato+ axons are not present in the thoracic cord (Figure 3F, F’, F”). Ratiometric quantification of percentage of axons at T1-T2 over C1-C2 (T:C ratio) shows that there is significantly more variability in the number of eGFP+ axons present in the thoracic dorsal funiculus than the number of tdTomato+ axons (Figure 3G). Note that in one mouse, ∼20% of eGFP+ axons present at C1-C2 reach thoracic T1-T2. In contrast, there is very little variation in the percentage of tdTomato+ axons that reach thoracic T1-T2 across 12 injected mice. These results highlight the relatively greater lack of specificity of labeling using AAV-eGFP injection into lateral cortex, which can still result in limited transduction of thoraco-lumbar-projecting CSN_TL_ in medial cortex. This indicates that Klhl14-Cre+ CSN extend axons only to the cervical spinal cord and not beyond, i.e. *Klhl14*+ CSN are cervical-projecting CSN (CSNc). These results confirm the specific expression of Cre by *Klhl14*+ CSN_BC-lat_ in Klhl14-Cre mice and highlight that this specificity enables selective targeting of CSN_BC-lat_ without solely relying on their spatial separation from CSN_med_ in the developing neocortex.

### Klhl14-Cre+ SCPN exhibit identical projections in the brainstem as all CSN_BC-lat_

Our results with both anterograde and retrograde labeling show that Klhl14-Cre is expressed by CSN_BC-lat_ and that Klhl14-Cre+ SCPN span both CBN and CSNc in lateral cortex. Since retrograde labeling does not ensure that all neurons will be labeled from a given location, we also used anterograde labeling to investigate whether Klhl14-Cre preferentially drives Cre expression in either CBN or CSN in lateral sensorimotor cortex. We therefore compared axonal collateralization of Klhl14-Cre+ SCPN in the brainstem with those of anatomically labeled CSN_BC-lat_. Our hypothesis was that if Klhl14-Cre labels a subset of CSN_BC-lat_ in lateral sensorimotor cortex, then we would find specific brainstem regions that would be innervated by anatomically labeled CSN_BC-lat_, but not by Klhl14-Cre+ SCPN. We therefore analyzed brainstem sections from P15 Klhl14-Cre mice that had received co-injections of AAV-eGFP and AAV-FLEX-tdTomato into lateral cortex at P0 (Figure 4A). We compared axonal collateralization by all CSN_BC-lat_ (eGFP+) with Klhl14-Cre+ SCPN (tdTomato+), spanning the entire rostro caudal extent of the brainstem from midbrain to caudal medulla (Figure 4B). Overall, we find very little difference between the two groups (Figure 4C-F). In a few areas in the medial midbrain, we find very subtle differences in labeling intensity between the two groups of axons; however, overall, both CSN_BC-lat_ and Klhl14-Cre+ SCPN axons display nearly identical patterns of collateralization across the entire rostrocaudal axis. These results indicate that Klhl14-Cre expression spans across the majority of SCPN in lateral sensorimotor cortex and reinforce the idea that Cre expression broadly occurs non-preferentially across all CSN_BC-lat_.

**Figure 4.**
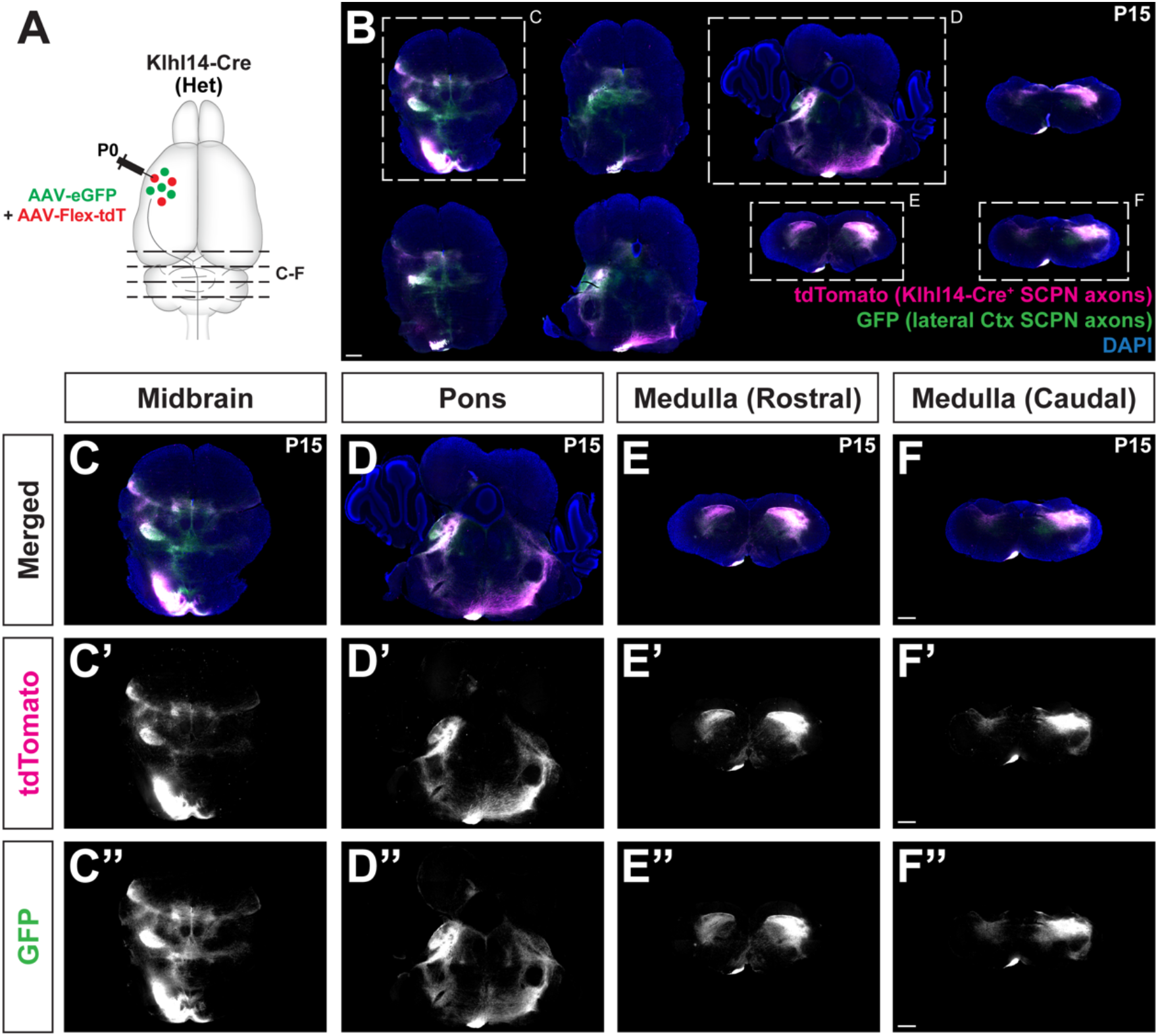
Klhl14+ SCPN axons exhibit identical projections in the brainstem as anatomically labeled CSN_BC-lat_. **A**, Similar experimental outline as in Figure 3. Klhl14-Cre knock-in mice were co-injected with AAV-eGFP and AAV-FLEX-tdTomato into rostrolateral cortex at P0. eGFP+ axons are from CSN_BC-lat_. The brainstem from injected mice was analyzed at P15. **B**, Coronal sections of the brainstem from one such injected mouse spanning the rostrocaudal extent of the brainstem showing axonal collateralization by Klhl14-Cre+ SCPN (magenta) vs. anatomically defined eGFP+ CSN_BC-lat_ (green). **C-F**, Magnified view of sections highlighted in B showing midbrain (C), pons (D), rostral medulla (E), and caudal medulla (F). Klhl14-Cre+ SCPN axons exhibit nearly identical collateralization across the brainstem as eGFP+ CSN_BC-lat_ axons. Scale bars are 500 µm.

### Misexpression of Crim1 and Cbln1 in Klhl14-T2A-Cre mice redirects Klhl14+ CSN axons to thoracic cord

In our prior work, we identified two novel regulators of CSN axon extension – Crim1 (Sahni et al., 2021b), and Cbln1 (Song *et al*., 2023)– that when misexpressed in CSN_BC-lat,_ can redirect these axons to extend past the cervical cord toward thoraco-lumbar segments. Since Klhl14-Cre provides a novel tool to selectively target CSN_BC-lat_, we next investigated whether Crim1 or Cbln1 misexpression can similarly re-direct Klhl14-Cre+ CSNc axons. We co-injected the lateral cortex of Klhl14-Cre mice at P0 with two AAVs: 1) AAV-FLEX-tdTomato to label CSN_BC-lat_ axons, and 2) one of the following three AAVs: either AAV-eGFP (control; 12 mice analyzed in Figure 3), AAV-Crim1, or AAV-eGFP-2A-Cbln1. We could not use eGFP in AAV-Crim1 due to the limited packaging capacity of AAVs (Figure 5A). We perfused injected mice at P15, and analyzed the spinal cords from all three groups.

**Figure 5.**
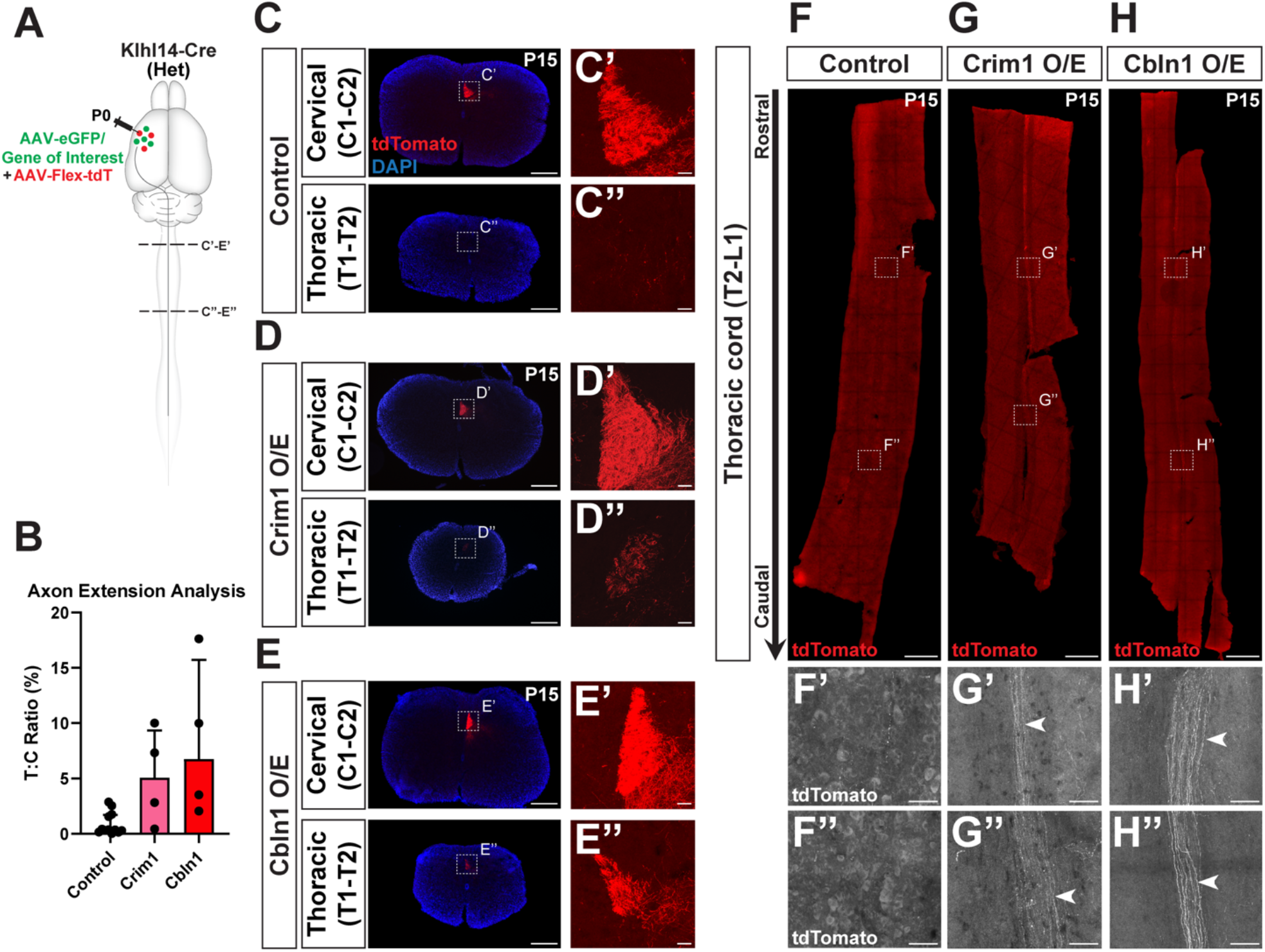
Misexpression of Crim1 and Cbln1 in Klhl14-T2A-Cre Mice Redirects Klhl14+ CSN Axons to Thoracic Cord. **A**, Schematic of experimental outline: P0 Klhl14-Cre heterozygous mice were co-injected with AAV-FLEX-tdTomato and either one of following three AAVs 1) AAV-eGFP (control); AAV-Crim1(Crim1 O/E) or AAV-Cbln1 (Cbln1 O/E) into rostrolateral cortex. **B**, Graph of average T:C ratios for controls (same results as in Figure 3G) (n=12) compared to Crim1 O/E (n=4) and Cbln1 O/E (n=4). Overexpression of either gene results in an increase in percentage of Klhl14-Cre+ CSN axons that extend into thoracic cord compared to control mice. **C-E**, Axial spinal sections from P15 Klhl14-Cre mice injected with either control (C-C”), Crim1 O/E (D-D”), or Cbln1 O/E (E-E”) at cervical and thoracic levels. Klhl14-Cre+ (tdTomato+) axons are present in the cervical dorsal funiculus (C’, D’, E’); however, only in Crim1 O/E (D”) and Cbln1 O/E (E”) mice do CSN axons extend to the thoracic dorsal funiculus. **F-H**, Maximum intensity projection images of thoracic cord horizontal sections (T2-T13) for control (F), Crim1 O/E (G), and Cbln1 O/E (H) groups. Monochrome magnified views of rostral and caudal levels from each cord are shown in F’-H”. In control mice, no CST axons are present within the first few thoracic segments (F’). In contrast, labeled tdTomato+ CSNc axons in both Crim1 O/E (G’) and Cbln1 O/E (H”) mice extend to far caudal levels in the thoracic cord (white arrowheads). Scale bars are 250 µm for **C, D, E**, 20 µm for insets in **C’-C”, D’-D”, E’-E”**, 500 µm for **F-H**, and 50 µm for **F’-F”, G’-G”, H’-H”**.

We first analyzed axial spinal sections at cervical C1-C2 and thoracic T1-T2 as in Figure 3. TdTomato+ (Klhl14-Cre+) CSNc axons in mice that received control AAV-GFP limit their extension to the cervical cord as previously described (Figures 5C-C”). In contrast, axial spinal sections from mice that were injected with either AAV-Crim1 or AAV-eGFP-2A-Cbln1, show substantial numbers of tdTomato+ (Klhl14-Cre+) axons extending to thoracic T1-T2 (Figures 5D-D”, E-E”). Quantification of these results (Figure 5B) confirmed that both *Crim1* and *Cbln1* can redirect *Klhl14*+ CSNc axons past their normal targets in the cervical cord toward the thoracic cord. We next analyzed the thoracic spinal cords from all three groups of injected mice. As expected, *Klhl14*+ CSNc axons in mice that received control AAV-GFP did not extend significantly into the thoracic cord (Figure 5F-F”). In striking contrast, horizontal sections of thoracic cord spanning thoracic T2-T13, from mice that received either AAV-Crim1 or AAV-eGFP-2A-Cbln1, show *Klhl14*+ CSNc axons extending to far caudal levels of thoracic cord (Figure 5G-H”). Together, these data indicate that known regulators of CSN axon extension that can re-direct CSN_BC-lat_ axons, are capable of similarly re-directing Klhl14-Cre+ CSNc axons.

### An Intersectional Approach to Target and Manipulate Klhl14+ SCPN

Our results thus far have identified that Cre expression in P0 Klhl14-Cre mice delineates CSN_BC-lat_ in lateral sensorimotor cortex, even though there is broader Cre expression in embryonic neocortex. We therefore wanted to investigate whether this expression specificity at P0 could be used to specifically target CSN_BC-lat_ at maturity. Our approach of using conditional retrograde labeling using rAAVs specifically labeled Klhl14+ SCPN in lateral sensorimotor cortex, i.e. CSN_BC-lat_; however, for circuit-level functional investigations of Klhl14+ SCPN, e.g., via silencing or activation experiments using AAV-DREADDs, it would be imperative that this method of labeling CSN_BC-lat_ does not also target other *Klhl14*+ cells in the nervous system. We therefore checked whether our conditional retrograde AAV labeling from the cerebral peduncle (as shown in Figure 2A) resulted in Cre-labeled neurons outside the cerebral cortex (schematized in Figure 6A). As shown previously, this strategy labels eGFP+ (Klhl14-Cre+) SCPN in lateral layer V (Figure 6B). We next analyzed coronal brainstem sections in these injected mice, i.e., near the rAAV injection site to see if we could find any eGFP+ neurons. We find widespread labeling of cell bodies throughout the rostro-caudal extent of the brainstem (Figures 6C-D”). These results indicate that while this approach of using conditional rAAV injections from the cerebral peduncles specifically labeled CSN_BC-lat_ in cortex, it also labels Klhl14-Cre+ neurons in the brainstem and therefore cannot be used to only target CSN_BC-lat_ in the CNS. Therefore, conditional Cre-dependent retrograde labeling from the cerebral peduncle at P0 in Klhl14-Cre mice cannot be used for circuit- and behavioral-level investigations of CSN_BC-lat_ function. To circumvent this problem, we next adopted a dual, intersectional viral labeling strategy to specifically label CSN_BC-lat_ without infecting Klhl14-Cre+ neurons in the brainstem.

**Figure 6.**
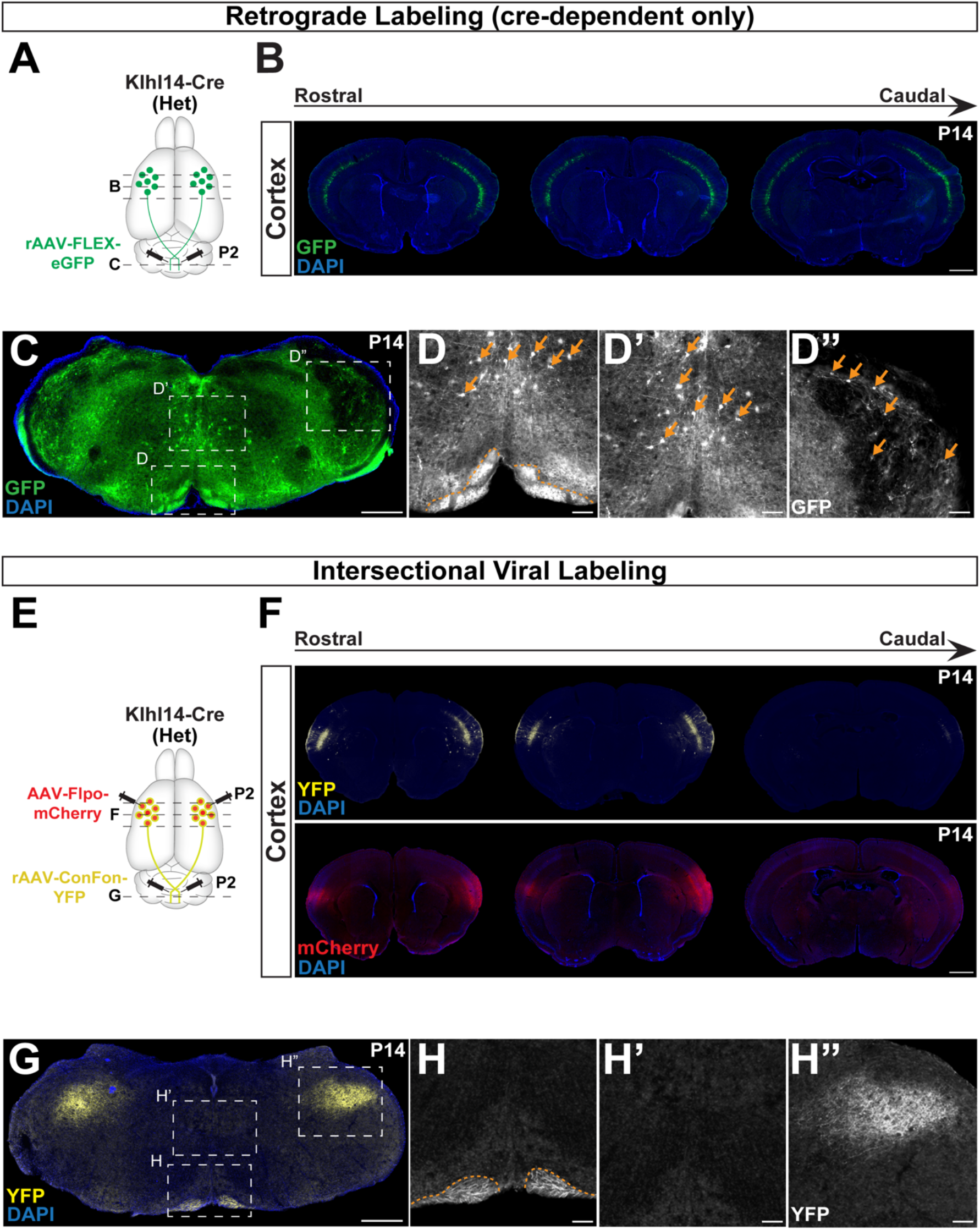
An Intersectional Approach to target Klhl14+ SCPN at maturity using Klhl14-Cre mice. **A**, Schematic of experimental outline: Klhl14-Cre mice received a bilateral retrograde injection of a retro AAV-FLEX-eGFP into the cerebral peduncle at P3. **B**, Coronal brain sections from an injected mouse at P14 show eGFP+ (Klhl14-Cre+) SCPN in lateral sensorimotor cortex. **C**, A representative section from the medulla from an injected mouse shows numerous eGFP+ neurons labeled at medial (D, D’) and lateral (D”) levels. Retrograde injection of AAV-FLEX-GFP therefore also labels Klhl14-Cre+ neurons in the brainstem. **E**, Schematic of experimental outline: Klhl14-Cre mice received bilateral intracortical injections of AAV-Flpo-mCherry into rostrolateral cortex at P2 along with bilateral retrograde injections of AAV-ConFon-eYFP in the cerebral peduncle. **F**, Coronal brain sections from a P14 injected mouse showing AAV-Flpo-mCherry (red) in lateral cortex. YFP+ (Cre+ and FlpO+) SCPN, i.e., Klhl14-Cre+ SCPN are specifically labeled in lateral cortex. (G) Representative coronal section of the medulla from the same mouse finds YFP+ (Klhl14-Cre+) SCPN axons (H), but no cell bodies are YFP+ (H’, H”). Therefore, this intersectional viral labeling approach labels Klhl14-Cre+ SCPN in cortex, but avoids labeling of Klhl14-Cre+ neurons in the brainstem, highlighting the effectiveness of this intersectional strategy for only targeting Klhl14-Cre+ SCPN at maturity. Scale bars are 1mm in **B, F**, 500 µm for **C, G**, and 100 µm for **D-D”, H-H”**.

We injected a rAAV-Cre-On-Flp-On-eYFP (ConFon-eYFP) (Fenno et al., 2014) into the cerebral peduncles at P2, followed by an anterograde AAV-Flpo-mCherry injection into lateral cortex (strategy schematized in Figure 6E). In this intersectional viral labeling approach, eYFP expression occurs only in neurons that express both Cre and FlpO. We next analyzed the cortex and brainstem from P15 Klhl14-Cre mice that had received these intersectional viral injections for eYFP labeled neurons. As expected, we find that CSN_BC-lat_ are eYFP+ in neocortex (Figure 6F); however, while Klhl14+ CSN_BC-lat_ axons are clearly eYFP+ in the ventral medulla (Figure 6G, H), we find no eYFP+ cell bodies in the brainstem (Figure 6H’). We similarly do detect CSN_BC-lat_ axon collaterals in the spinal trigeminal nucleus but no neuronal labeling in this region (compare Figure 6H” with 6D”). Therefore, this intersectional labeling approach enabled selective targeting of only CSN_BC-lat_ in the CNS without labeling Klhl14-Cre+ neurons in the brainstem. Our results show that Klhl14-Cre mice are a novel tool for the prospective identification and investigation of CSN_BC-lat_ axon targeting during development. In addition, intersectional viral labeling can expand their utility for functional analyses of CSN_BC-lat_ in the mature CNS.

## Discussion

The CST is a principal circuit for skilled voluntary motor control, which necessitates that SCPN axons make segmentally specific connectivity with their subcerebral targets. Our previous work had identified molecular delineation between developing SCPN subpopulations in sensorimotor cortex that will eventually extend axons to different levels of the neuraxis. *Crim1* expression could prospectively identify CSN_TL_, which suggested that this principle could be applicable to other subpopulations. In the present study, we establish that *Klhl14* expression, at specific developmental times, can prospectively delineate another CSN subpopulation – CSN_BC-lat_. These new results establish that developmental molecular delineation establishes a first step towards durable specificity of corticospinal connectivity at maturity. We further establish that Khl14-Cre provides a novel tool to reproducibly and reliably investigate CSN_BC-lat_ axon targeting with greater precision and specificity. This presents a significant advance over previous approaches that were reliant on anatomical separation of this subpopulation within sensorimotor cortex. Finally, we establish that, using intersectional viral labeling, Khl14-Cre can be used as a novel tool to specifically target CSN_BC-lat_ in the adult CNS. Our results indicate that molecular delineation early in development can be used to investigate corticospinal subpopulation-specific development, axon extension, connectivity, and eventually, function at maturity.

What makes Klhl14 a highly desirable tool to investigate SCPN subpopulation development and connectivity is that it is expressed with high specificity during development and is not expressed in adulthood. Several genes that exhibit highly specific expression during development are either absent in the mature cortex, or even worse, become expressed more broadly in the adult brain. As a result, in such instances, adult gene expression cannot be used for targeting specific neuronal populations in the adult brain. Therefore, using developmental gene expression provides a valuable tool to target cell populations of interest, provided they can be accessed at the appropriate time in development. While inducible Cre lines do provide the ability for such temporal control, our approach shows that developmental anatomical labeling in Cre mouse lines can provide exquisite specificity for investigation and potential manipulation of SCPN subpopulations.

Cre expression in Klhl14-Cre mice recapitulated our previously established specificity of *Klhl14* expression by CSN_BC-lat_; however, this specificity occurs at a specific developmental time. Interestingly, our results using genomic reporters find much broader Cre-dependent recombination indicating widespread Cre expression early in development, suggesting a potentially earlier role for Klhl14 in neocortical development. While it remains theoretically possible that this earlier Cre expression is driven by regulatory elements that are not associated with the Klhl14 locus, the specificity of expression at P0 suggests that this is likely not the case. Regardless, the approach of conditional labeling using AAV-mediated gene delivery enabled investigation of CSN_BC-lat_ with specificity. This also suggests that a similar principle could apply to other Cre lines that, when crossed to genomic reporters of Cre-dependent recombination, display broader expression that could be wrongly construed as non-specific. A vast repertoire of Cre lines have now been made available (Daigle et al., 2018; Gerfen et al., 2013) and while there remain legitimate concerns regarding the specificity of expression (Luo et al., 2020; Stuber et al., 2015) and/or leaky transgene expression when using Cre-dependent labeling via AAVs (Botterill et al., 2021), our results indicate that discrepancy between results from such reporters and “established” expression data could also arise from spatiotemporal dynamics of gene expression. To conclusively establish gene expression requires integration across multiple approaches that can both qualitatively and quantitatively determine where and at what point in development a specific gene is expressed. Molecular strategies that integrate this critical information regarding gene expression dynamics across spatiotemporal dimensions are likely to be more precise and tailored for gene targeting and functional manipulation experiments.

Toward this goal, large-scale sequencing datasets are now becoming rapidly available, providing a valuable resource for using gene expression dynamics to gain molecular access to defined neuronal populations (Di Bella et al., 2021; Rosenberg et al., 2018; Tasic, 2018; Tasic et al., 2016; Yao et al., 2021). These datasets are beginning to provide greater resolution of such dynamics across the developing and adult CNS and presumably will provide greater insights into the necessary subtleties of gene expression that can be utilized to target specific cell types with precision. In line with this, in more recent experiments, we used single cell transcriptomics (scRNA-seq) to investigate additional diversity within CSN_BC-lat_. Our results are that CSN_BC-lat_ comprises additional diversity, with both cortico-brainstem and cervical-projecting corticospinal neurons residing interdigitated in lateral sensorimotor cortex (Kaiser et al., 2022). ScRNA-seq finds that while *Klhl14* is expressed by both cortico-brainstem and corticospinal subsets, cervical-projecting corticospinal neurons express Klhl14 at a lower level (Kaiser *et al*., 2022). Consistent with this, our results using Klhl14-Cre mice highlight that *Klhl14* is expressed by both populations in lateral sensorimotor cortex, with a subset of Klhl14+ SCPN extending axons to the cervical spinal cord. Since retrograde labeling always has the caveat that it is impossible to label all neurons from any given level of the neuraxis, we used anterograde labeling, which is more sensitive, to investigate the extent of coverage of CSN_BC-lat_ by Klhl14-Cre. Analysis of axonal collateralization by Klhl14+ SCPN axons across the entire rostro caudal extent of the brainstem indicates that Klhl14-Cre broadly labels CSN_BC-lat_ with extensive innervation across the brainstem and cervical spinal cord. Klhl14-Cre does not differentiate between cortico-brainstem versus corticospinal neurons in lateral sensorimotor cortex, which we recently identified to be molecularly distinct (Kaiser et al. 2022).

Our quantitative data reveal that Klhl14-Cre+ SCPN early in development largely comprise cortico-brainstem neurons and a smaller subset of spinal projecting neurons. scRNAseq results show that corticospinal neurons in lateral cortex express lower levels of *Klhl14*, and therefore, it is possible that this is why we detect fewer Klhl14-Cre+ corticospinal neurons. It is, however, interesting to note that Klhl14+ SCPN includes a population of cervical projecting corticospinal neurons even at maturity. It is well established that SCPN axons exhibit extensive pruning throughout development, such that several SCPN eventually lose their spinal projections (Abe et al., 2024; Stanfield and O’Leary, 1985; Stanfield et al., 1982). Therefore, significant numbers of anatomically labeled corticospinal neurons early in development can differentiate into cortico-brainstem neurons at maturity. Our results reveal that Klhl14-Cre+ corticospinal neurons maintain their spinal projection past this developmental period of axon pruning and therefore remain corticospinal neurons even at maturity.

*Klhl14* is only transiently expressed by SCPN during development, and single cell sequencing datasets find that *Klhl14* expression is largely absent in adult cortex(Yao et al., 2021; Yao et al., 2023). Using retrograde labeling in Klhl14-Cre mice, we were therefore able to map the areal locations of Klhl14-Cre+ SCPN and CSN during development, at maturity. Consistent with the predictions from our previous developmental expression analyses, we find that the overwhelming majority of Klhl14-Cre+ SCPN and CSN reside in somatosensory cortex. While a small minority of Klhl14+ SCPN are annotated as located in primary motor cortex, these neurons actually reside in the transition zone between agranular motor cortex and granular somatosensory cortex (Tennant et al., 2011).

Since *Klhl14* expression in absent in adulthood, our results cannot establish whether Klhl14-Cre+ SCPN maintain molecular distinctions from other SCPN subpopulations at maturity. However, recent single nuclear profiling across all descending spinal projecting neurons in the mouse adult brain finds that spinal projecting neurons within primary somatosensory and motor cortex are comparatively similar (Winter et al., 2023). This suggests that while Klhl14-Cre expression during development primarily delineates corticospinal projections arising from primary somatosensory cortex (Figure 2D), these developmental molecular differences are not maintained into maturity. Performing longitudinal molecular profiling of Klhl14-Cre+ SCPN through development into adulthood might identify potentially more subtle differences that could differentiate them from other SCPN subpopulations in the adult sensorimotor cortex. More importantly, these results highlight the fact that developmental molecular controls over such differentiation are only expressed at the specific developmental time and that these molecular differences are largely not maintained into maturity.

We also established that Klhl14-Cre mice enable specificity of targeting CSN_BC-lat_ for analyzing axon targeting in the spinal cord. Even in instances where the AAV injection spread beyond the injection site in lateral cortex, axon targeting analysis showed that Klhl14-Cre+ axons remained restricted to the cervical cord. This specificity also enabled investigating molecular control over CSN_BC-lat_ axon targeting in the spinal cord. Consistent with the fact that Klhl14-Cre expression labels all CSN_BC-lat_, we find that misexpression of either of previously identified molecular regulators, *Crim1* and *Cbln1*, can redirect *Klhl14*+ CSN_BC-lat_ axons in a similar manner. Since CSN_BC-lat_ reside in lateral sensorimotor cortex, the strategy in previous experiments was aimed at restricting misexpression in CSN_BC-lat_ by injecting AAVs into lateral, but not medial, sensorimotor cortex. (Sahni et al., 2021a; Sahni et al., 2021b; Song et al., 2023). Since AAVs are known to spread from the injection site, AAV injected into lateral cortex could potentially spread to medial cortex. This is especially likely in the developing cortex, where the spatial separation between these regions is smaller than in the adult. As a result, specifically targeting AAVs to CSN_BC-lat_ using anatomical separation alone is experimentally challenging.

Consequently, in previous experiments, any mice in which AAV injections spread into medial cortex, had to be excluded from analysis, because in these mice, the labeled axons were no longer exclusively CSN_BC-lat_. In contrast, Klhl14-T2A-Cre mice enable selective labeling of CSN_BC-lat_ making it possible to investigate CSN_BC-lat_ axon targeting even in instances where AAV injections spread into medial cortex. This therefore provides a significant advance over our previous strategy and will enable testing function of other genes in a relatively more high-throughput manner.

In addition, our previous work identified Klhl14 as likely functioning as a transcriptional repressor, whereby it represses expression of both Crim1 and Cbln1 in CSN_BC-lat_, which would otherwise direct these axons to caudal spinal segments (Sahni et al., 2021b) (Song *et al*., 2023). Our results with Klhl14-Cre mice now conclusively establish that *Klhl14*+ SCPN axons are responsive to misexpression of either of these genes. These new results further validate and confirm these previously identified genetic interactions. This establishes that Klhl14 functions in CSN_BC-lat_ to represses CSN_TL_ gene expression and restrict axonal projections to proximal targets in the brainstem and spinal cord.

Our results lay the foundation for establishing and using novel Cre-mouse lines for targeting and investigating specific CSN subpopulations through development into maturity. Such tools can then be used to analyze the molecular control over their development, their anatomical organization at maturity, and potentially specific, circuit-level functional roles in adulthood. The CST is a critical circuit that has been investigated for functional recovery in several instances of nervous system damage. Collectively, results from such CSN-subpopulation-specific targeting and analyses can begin to elucidate mechanisms that not only control normal CST development, but that can also be utilized to effect regeneration and plasticity in instances of disease and injury. Our results are beginning to lay the foundation for such investigations into links between CST development and adult CST repair.

